# Impact of admixture and ancestry on eQTL analysis and GWAS colocalization in GTEx

**DOI:** 10.1101/836825

**Authors:** Nicole R. Gay, Michael Gloudemans, Margaret L. Antonio, Brunilda Balliu, YoSon Park, Alicia R. Martin, Shaila Musharoff, Abhiram Rao, François Aguet, Alvaro Barbeira, Rodrigo Bonazzola, Farhad Hormozdiari, GTEx Consortium, Kristin G. Ardlie, Christopher D. Brown, Hae Kyung Im, Tuuli Lappalainen, Xiaoquan Wen, Stephen B. Montgomery

## Abstract

**Background:** Population structure among study subjects may confound genetic association studies, and lack of proper correction can lead to spurious findings. The Genotype-Tissue Expression (GTEx) project largely contains individuals of European ancestry, but the final release (v8) also includes up to 15% of individuals of non-European ancestry. Assessing ancestry-based adjustments in GTEx provides an opportunity to improve portability of this research across populations and to further measure the impact of population structure on GWAS colocalization.

**Results:** Here, we identify a subset of 117 individuals in GTEx (v8) with a high degree of population admixture and estimate genome-wide local ancestry. We perform genome-wide *cis*-eQTL mapping using admixed samples in six tissues, adjusted by either global or local ancestry. Consistent with previous work, we observe improved power with local ancestry adjustment. At loci where the two adjustments produce different lead variants, we observe only 0.8% of tests with GWAS colocalization posterior probabilities that change by 10% or more. Notably, both adjustments produce similar numbers of significant colocalizations. Finally, we identify a small subset of GTEx v8 eQTL-associated variants highly correlated with local ancestry (R^2^ > 0.7), providing a resource to enhance functional follow-up.

**Conclusions:** We provide a local ancestry map for admixed individuals in the final GTEx release and describe the impact of ancestry and admixture on gene expression, eQTLs, and GWAS colocalization. While the majority of results are concordant between local and global ancestry-based adjustments, we identify distinct advantages and disadvantages to each approach.

## Background

Thousands of genome-wide association studies (GWAS) have been published to date. Subsequently, large-scale expression quantitative trait loci (eQTL) datasets are studied to provide insights for genetic variants associated with complex traits. While the majority of such studies focus on single-ancestry populations or relatively homogeneous populations, the latest Genotype-Tissue Expression (GTEx) project (v8) includes up to 17% of individuals with non-European or admixed ancestry (1). Genetic studies with individuals of admixed ancestries may suffer from additional challenges due to complex population substructure (2,3). Such substructure can lead to confounding genetic associations, and insufficient control may increase spurious findings (4,5).

Global ancestry (GA), or the proportion of different ancestral populations represented across the entire genome, is routinely used to adjust for population structure in genetic association studies (6). This approach has the advantage of averaging genomic background effects and was used in eQTL mapping for the main GTEx releases (1,7). The potential disadvantage of correcting only for GA is that it does not precisely account for ancestry at any specific locus. The can be problematic when genes are differentially expressed in ancestral populations of admixed individuals. In contrast, local ancestry (LA), or the number of alleles derived from distinct ancestral populations at a given locus, may be more appropriate for population structure adjustment in admixed populations but typically suffers from much longer compute time and can be prone to errors in estimation at a variant level (5,8–12).

LA adjustment in genetic association studies has been shown to reduce type I error rate (false positives) (13–15) and sufficiently control for population stratification (13,15). However, the power of adjusting for LA is highly dependent on the underlying genetic architecture of the admixed population (8,12,15–17); some have recommended using LA adjustment as a method for follow-up of candidate loci as opposed to a discovery tool for GWAS (8,14,18). Fewer studies have investigated the effect of LA adjustment on eQTL mapping, demonstrating modest improvements in discovery power (5,10). Recently, Zhong *et al*. have demonstrated that the use of LA adjustment, compared to GA adjustment, can improve eQTL mapping while controlling for type I error rate and increasing statistical power (10). However, the implications of these differences for GWAS colocalization were not assessed.

In this study, we describe the degree of admixture in the GTEx v8 cohort and estimate LA for a subset of 117 individuals with at least 10% admixture from European, African, and Asian ancestral populations. LA explains at least 10% of variance in residual expression for 1% of expressed genes (N=183). We perform *cis*-eQTL mapping in six tissues and assess the differences between LA adjustment and GA adjustment in the context of this admixed subcohort. For the subset of loci where the two ancestry adjustment methods yield different results, we perform GWAS/eQTL colocalization analyses with 114 previously published GWAS. We assess the effect of within-continent and between-continent population stratification of the eQTL-associated variants (eVariants) on differences in colocalization between the two ancestry adjustment methods. Finally, we identify a small subset of GTEx eVariants whose genotypes are highly correlated with LA, providing a resource to enhance functional follow-up of these loci.

## Results

### The final GTEx release includes African and Asian population admixture

The GTEx v8 release includes whole genome sequencing and gene expression data for 838 individuals, including 103 African American and 12 Asian American individuals (selfreported ancestry). Genome-wide genotype-based principal components (gPCs) reflect GA and have been used to adjust for population structure in both GWAS (6,9,13) and eQTL studies (7).

Therefore, to understand the degree of population admixture represented in GTEx, we compared the first two gPCs with self-reported ancestry (Figure 1a). Figure 1a demonstrates that gPC1 and gPC2 reflect African and Asian ancestry, respectively; the majority of European Americans (698 out of 714 individuals) cluster together near the origin, suggesting that the samples in this cluster are relatively homogeneously European-descendent. These patterns are observed with finer resolution when genotype PCA is performed with combined GTEx and 1000 Genomes data (Figure S1, Additional file 1). A subset of 117 individuals with more than 10% population admixture, referred to as 117AX, were retained for downstream analyses (Figure 1a; Table S1, Additional file 2).

**Figure 1.**
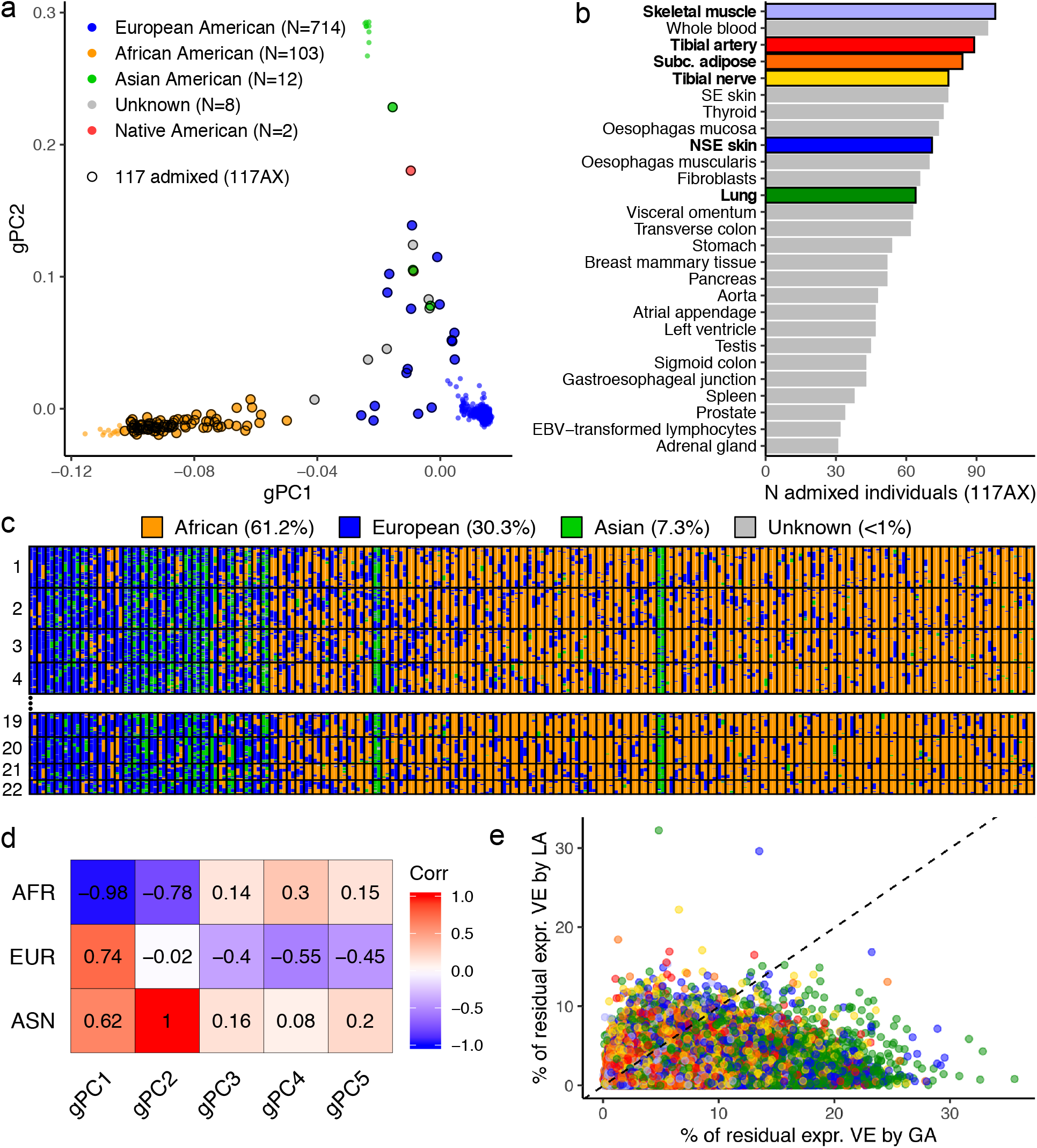
Population admixture in the GTEx v8 cohort. **(a)** Genotype principal components (gPCs) reflect global ancestry. Points are colored by selfreported ancestry. Circled points indicate the 117 individuals defined as admixed (117AX). **(b)** A subset of GTEx v8 tissues have an 117AX sample size of at least 30. The six tissues selected for *cis*-eQTL mapping in 117AX are colored and shown in bold. **(c)** LA tracts collapse consecutive variants on a single parental chromosome with the same ancestry assignment into contiguous haplotype blocks. The fine spatial resolution of local ancestry contrasts with the global ancestry proportions indicated in the legend. Haplotypes (columns) are paired by individual; rows are autosomal chromosomes. Individuals are sorted from left to right by decreasing proportions of European admixture. **(d)** gPCs are highly correlated with global ancestry proportions averaged from genome-wide local ancestry. **(e)** Local (or global) ancestry explains a fraction of variance in residual gene expression after correcting for global (or local) ancestry. Local ancestry is defined as the local ancestry at the transcription start site of each gene; global ancestry is the first five gPCs. Points are colored by tissue; colors correspond with **(a)**. VE = variance explained.

The 49 tissues used for QTL discovery in the GTEx v8 release have varying representation of 117AX. 27 of these tissues have a sample size of at least 30 admixed individuals (Figure 1b). Sample sizes for all 49 tissues are provided in Figure S2 (Additional file 1). The pituitary and 13 central nervous system tissues have the lowest representation of 117AX relative to total sample sizes per tissue (mean 7%). We selected six tissues in which to perform *cis*-eQTL calling based on a minimum admixed sample size of 60 (19) and well-studied trait relevance (20–24): subcutaneous (subc.) adipose (N=84), tibial artery (N=89), lung (N=64), skeletal muscle (N=98), tibial nerve (N=78), and not sun-exposed (NSE) skin (N=71).

Using RFMix (25), we performed three-population (European, African, and Asian) LA estimation on 117AX (see Methods; Figure 1c; Figure S3, Additional file 1). We provide these LA calls as a resource for further investigation of GTEx data (Table S2, Additional file 2). For each individual, genome-wide LA was averaged to provide GA estimates. Every sample in 117AX has less than 90% GA from any one ancestral population out of Europe, Africa, and Asia. We correlated these GA proportions with the first five gPCs, which quantitatively demonstrates the strong relationships between gPC1 and African ancestry (r = −0.98) and gPC2 and Asian ancestry (r = 1.0; Figure 1d).

In order to assess the importance of LA in the context of gene expression, we adapted an existing approach (26) to calculate the proportion of variance explained in 117AX gene expression by LA after accounting for GA and vice versa (see Methods; Figure 1e; Table S3, Additional file 2). On average, across genes in our six tissues of interest, GA explains more variance in gene expression than LA at the transcription start site for each gene (*P*-value < 2.2e-16, two-sided t-test). However, LA explains at least 10% of variance in residual expression for 1% of expressed genes (N=183). At the extreme, LA explains 32% of variance in residualized expression of *TBC1 Domain Family Member 3 (TBC1D3)*, a hominoid-specific oncogene (27), in Tibial Artery; LA also explains significantly more variance in *TBC1D3* expression than GA in all six tissues tested (two-sided t-test; *P*-value = 0.008). In a separate study of copy number, *TBC1D3* was among the most variable (median 38.13, variance 93.2 copies among 159 individuals) and population-stratified (mean 29.28, 34.17, and 43.86 copy numbers in European, Asian, and Yoruban samples, respectively) human gene families (28). Such biological evidence for residual variance in gene expression captured by LA supports the importance of considering LA in the context of eQTL mapping.

### Local ancestry adjustment increases power for discovery in *cis*-eQTL mapping

We performed *cis*-eQTL mapping in the admixed population (117AX) to identify associations between variants and gene expression within each of the six tissues indicated in Figure 1b (see Methods; Table S4, Additional file 2). We implemented linear models to test for an association between each gene-*cis*-variant pair. For each pair, two association tests were performed: the first to adjust for global ancestry (GlobalAA); the second to adjust for local ancestry (LocalAA). Importantly, LocalAA accounts for the number of European, African, and Asian alleles for each variant while GlobalAA uses the first five genotype principal components as a proxy for global ancestry, implementing the same ancestry adjustment used in the GTEx eQTL calling pipeline.

*A* quantile-quantile plot of the nominal *P*-values (-log10) of all association tests in GlobalAA and LocalAA demonstrates that LocalAA has more significant *P*-values (represented in the highest quantiles) relative to GlobalAA for five of the six tissues, with NSE skin showing more similar *P*-value distributions between the two methods (Figure 2a). This corroborates previous findings that LA adjustment results in more significant nominal *P*-values than GA adjustment in the context of cis-eQTL mapping (10).

**Figure 2.**
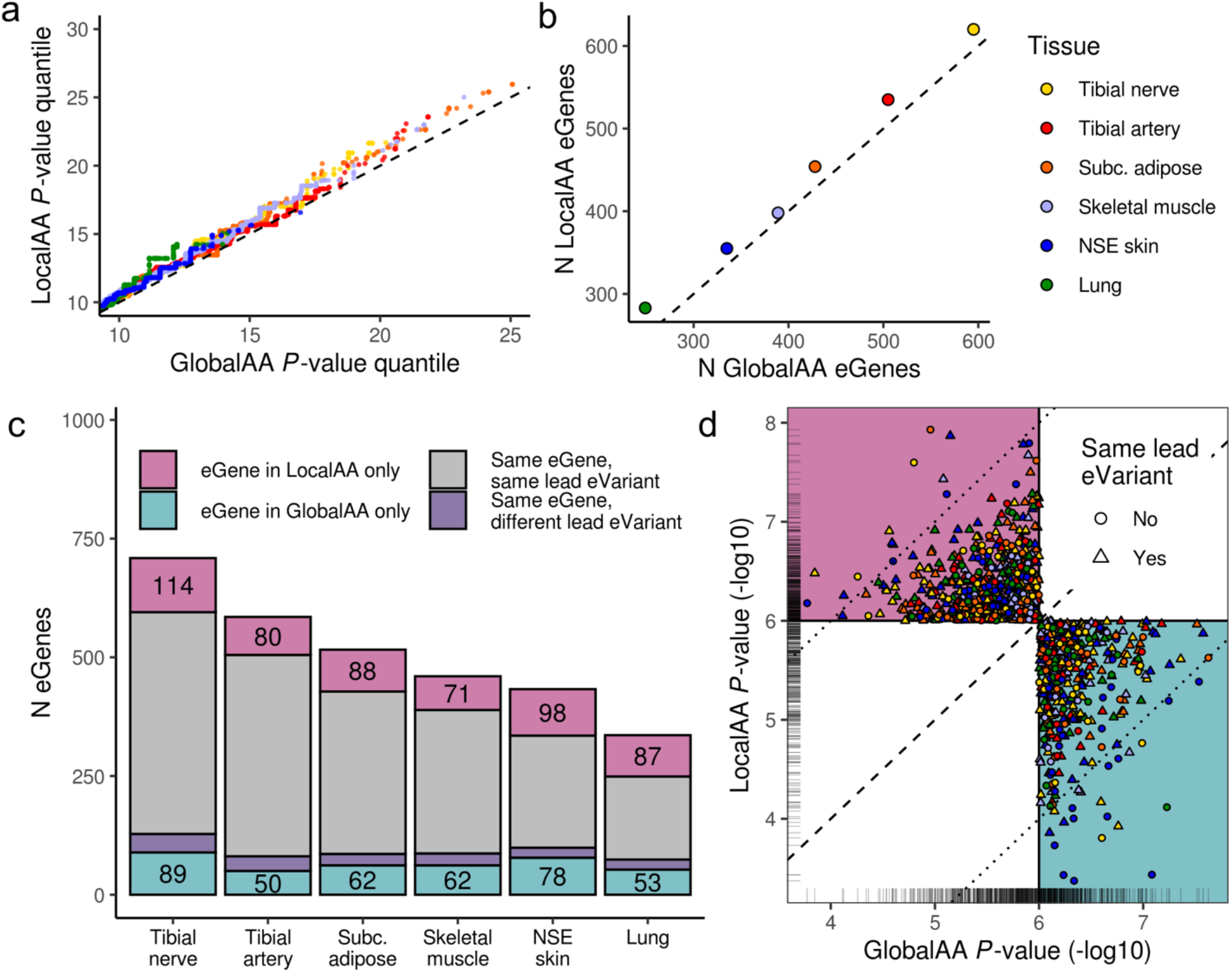
Comparison of *cis*-eQTLs called by LocalAA or GlobalAA. *Cis*-eQTL mapping was performed in six tissues. A nominal *P*-value threshold of 1e-6 was applied to identify significant associations. **(a)** A Q-Q plot of nominal *P*-values for all tests indicates a modest improvement of power in most tissues when using LocalAA. **(b)** LocalAA identifies more eGenes than GlobalAA in all six tissues (*P*-value=0.016, binomial probability). **(c)** The majority of eGenes are identified by both ancestry adjustment methods (gray + purple). The two methods report different eVariants for a small fraction of these eGenes (purple). Numbers indicate eGenes uniquely called by one of the ancestry adjustment methods, which are plotted in **(d). (d)** The majority of eGenes unique to one ancestry adjustment method fall near the significance threshold, as indicated by the rug plot. Dotted lines demarcate the region outside of which eGenes in one method have a nominal *P*-value at least two orders more significant than the alternate method. Points are colored by tissue.

We applied a nominal *P*-value cutoff of 1e-6 to identify significant eQTLs; this threshold closely approximates the threshold required for an eQTL to subsequently pass a false discovery rate cutoff of 5%. More eGenes are called with LocalAA than GlobalAA in all six tissues (*P*-value=0.016, binomial probability) (Figure 2b). The majority of eGenes overlap between the two methods, a subset of which have different associated lead eVariants between LocalAA and GlobalAA (Figure 2c). This subset of eGenes provided an opportunity to characterize differences in lead eVariants identified between the two ancestry adjustment methods and was the focus of downstream analyses.

eGenes are considered unique to an ancestry adjustment method if the association reaches significance only with that method (nominal *P*-value cutoff of 1e-6; 839 total instances across tissues for 794 unique genes). The majority (64%) of eGenes that are unique to one method replicate at a *P*-value within one order of magnitude of the other method (Figure 2d). However, 29 of these eGenes only replicate in the other method at a *P*-value more than two orders of magnitude less significant (10 and 19 eGenes unique to LocalAA and GlobalAA, respectively). 15 of these 29 eGenes are in NSE skin; none are in tibial artery. Interestingly, in 20 out of these 29 eGenes, despite the large difference in statistical significance, the lead variants between the two adjustment methods are identical.

### Different eQTL ancestry adjustments yield minor differences in GWAS colocalization

Colocalization analyses assess the degree to which independent signals of association share the same causal variant (posterior probability of colocalization 4, PP4). We performed colocalization between our twelve sets of eQTL summary statistics (one per ancestry adjustment method per six tissues) and 114 GWAS using COLOC (29). We restricted colocalization tests to loci surrounding the subset of eGenes with different lead eVariants between LocalAA and GlobalAA (Figure 2c; Table S5, Additional file 2). We hypothesized that improvements in eQTL analysis by either LocalAA or GlobalAA would be reflected in systematically higher posterior probabilities for eQTL/GWAS colocalization.

While GWAS colocalization was only tested at loci for which the two eQTL ancestry adjustment methods yielded different lead eVariants, colocalization probabilities are not systematically different between the two methods (*P*-value = 0.64, two-sided t-test); only 0.8% of tests have an absolute difference in PP4 greater than 0.1 (Figure 3a). Furthermore, loci with strong evidence of colocalization (PP4 > 0.5) have similarly high posterior probabilities of colocalization regardless of correction methods, indicating that robust effects are captured by both ancestry adjustments. Colocalizations are considered stronger with one ancestry adjustment method if two conditions are met: 1) PP4 > 0.5 for only one of the two eQTL adjustment methods; and 2) the absolute difference between GlobalAA PP4 and LocalAA PP4 is greater than 0.3. 23 loci meet these conditions (see Figure S4, Additional file 1).

**Figure 3.**
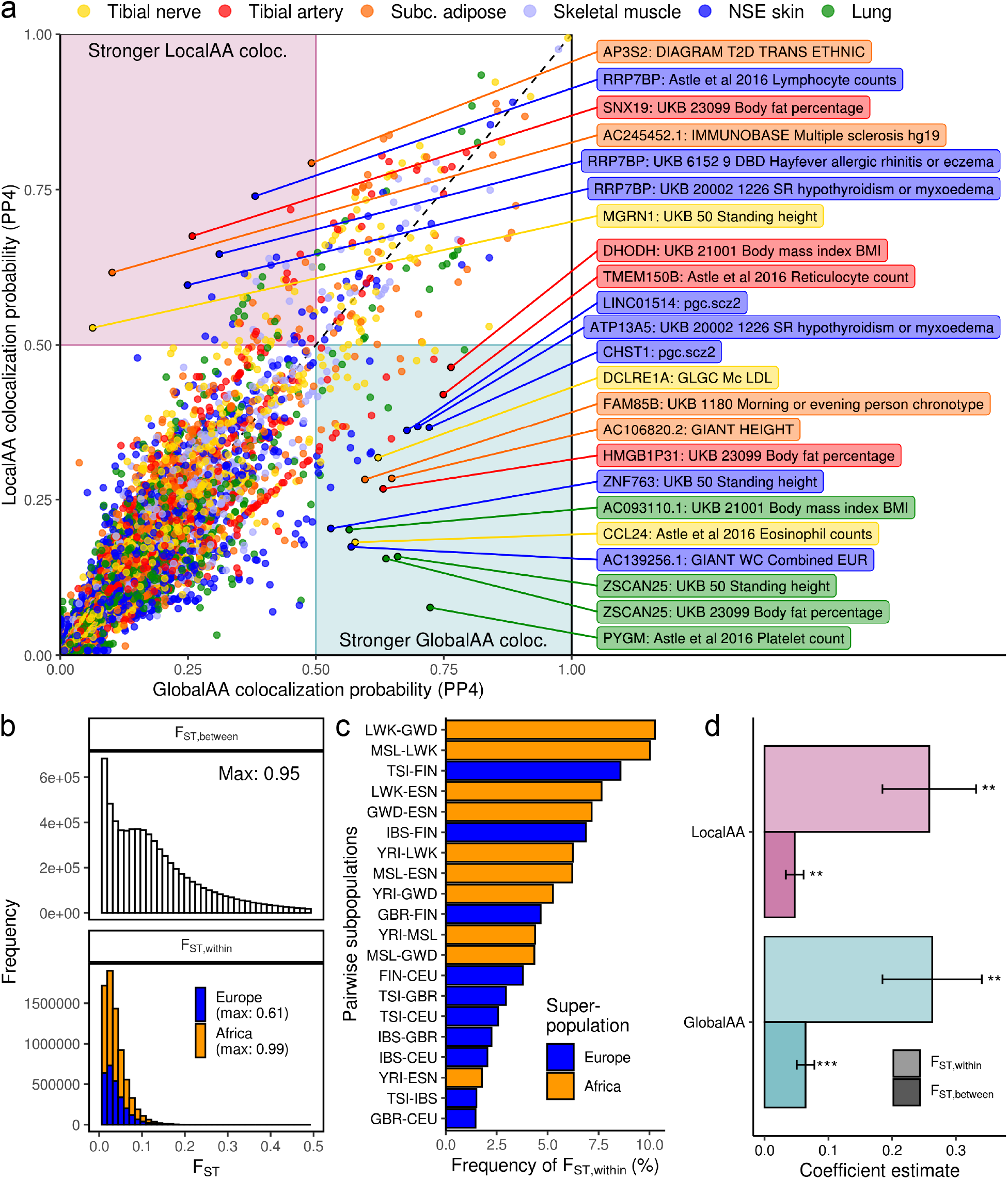
Impact of eQTL ancestry adjustment methods on colocalization with GWAS. **(a)** Each point represents a GWAS/eQTL colocalization test near a single eGene (colored by eQTL tissue). The y- and x-axes respectively show the posterior probability of colocalization (PP4) for this test using either LocalAA or GlobalAA eQTL statistics. We performed colocalization for the subset of loci where LocalAA and GlobalAA called eQTLs with different lead eVariants (nominal *P*-value threshold of 1e-4). Points are labelled with eGenes and GWAS traits if the colocalization is stronger with one eQTL ancestry adjustment method (see main text). SR = self-reported; DBD = diagnosed by doctor. **(b-d)** In order to identify the effect of population structure on PP4, we calculated within- and between-continent divergence in allele frequencies (F_ST_) for all variants tested during eQTL mapping. We used continental African and European 1000 Genomes subpopulations for these calculations. (b) Between-continent F_ST_ (F_ST,between_) is the F_ST_ between Africa and Europe. Within-continent F_ST_ (F_ST,within_) is the maximum of 20 F_ST_ calculated for pairwise subpopulations within Europe or Africa. (c) provides the frequency in which each of the 20 pairwise comparisons per variant yields F_ST,within_. (d) We tested the effects of F_ST,within_ and F_ST,between_ of lead eVariants on PP4. For each eQTL adjustment method, the best colocalization (maximum PP4) per eGene per tissue was identified; we regressed the maximum PP4 on F_ST,within_ and F_ST,between_ of the method’s lead eVariant. A positive regression coefficient indicates that larger values of the predictor (F_ST_) are associated with higher maximum probabilities of colocalization. Error bars indicate standard errors of the coefficient estimates. ** and *** indicate *P*-value < 1e-3 and *P*-value < 1e-5, respectively.

For the 23 loci where one adjustment method has a stronger colocalization, most colocalization probabilities are stronger for GlobalAA versus LocalAA (N = 16). However, the locus with the strongest colocalization probability favors colocalization with LocalAA (LocalAA PP4 = 0.79; GlobalAA PP4 = 0.49). There is evidence that this colocalization between the LocalAA *AP3S2* eQTL in subcutaneous adipose and the DIAGRAM Consortium Type II Diabetes (T2D) GWAS is biologically relevant (30). Variants near *AP3S2* have previously been associated with T2D in at least 8 other GWAS, one of which was included in the DIAGRAM meta-analysis (31). Many of these GWAS were performed in diverse or non-European populations (32–37). The eQTL tissue in this colocalization, subcutaneous adipose, also reflects known adipose pathologies associated with T2D (38). However, overall, we observe that neither LocalAA nor GlobalAA performs significantly better in the context of colocalization.

### Within- and between-continent genetic differences impact colocalization

We tested if population differentiation in LocalAA or GlobalAA lead eVariants affects colocalization probabilities. Fixation index, or F_ST_, is a measure of genetic population differentiation, where higher F_ST_ indicates more divergent allele frequencies between two populations (2,39). We calculated F_ST,within_ and F_ST,between_ for each variant tested during eQTL calling using 1000 Genomes populations and allele frequencies (see Methods; Figure 3b). We only included African and European populations in this analysis due to the relatively small representation of Asian ancestry in 117AX (7.3%, Figure 1c).

To define F_ST,within_ for a variant, we calculated F_ST_ for 10 pairs of European subpopulations and 10 pairs of African subpopulations; F_ST,within_ was defined as the maximum of these 20 values. As expected, F_ST_ is generally higher within Africa than within Europe, reflected in the higher number of variants with F_ST,within_ derived from a pairwise African population comparison (Figure 3c) (40). For example, 10% of tested variants have a maximum F_ST_ between LWK (Luhya in Webuye, Kenya) and GWD (Gambian in Western Divisions in the Gambia) out of the 20 pairwise subpopulation comparisons.

Linear regression was used to test the association between colocalization probability and F_ST,within_ and F_ST,between_ of lead eVariants. This regression was performed independently for each ancestry adjustment method; observations were limited to the strongest colocalization (maximum PP4) per eGene per tissue. A positive coefficient indicates that higher values of the predictor (F_ST_) are associated with stronger colocalization probabilities (maximum PP4). Both F_ST,within_ and F_ST,between_ are significant predictors of maximum PP4 for both ancestry adjustment methods (Figure 3d). Interestingly, F_ST,within_ has a larger coefficient than F_ST,between_ in both regressions. In other words, within-continent population differentiation at the lead eVariant has a greater impact on colocalization than between-continent population differentiation at the lead eVariant, regardless of the eQTL ancestry adjustment method.

Furthermore, the coefficients for FST,within are the same between GlobalAA and LocalAA. If GlobalAA and LocalAA colocalizations are similarly affected by within-continent population differentiation, then both ancestry adjustment methods may correct for within-continent population structure to similar degrees. This similarity suggests both LocalAA and GlobalAA may benefit by accounting for finer-scale reference populations or inclusion of more gPCs but does not exclude the potential that some high F_ST_ variants are also enriched for functional effects.

### A subset of GTEx v8 eVariants are highly correlated with local ancestry

One justification for performing LocalAA as opposed to GlobalAA is the unique ability to avoid confounding by local population structure (15). We examined the reported GTEx v8 significant associations for evidence of confounding with LA. For each eVariant in the set of all significant associations across tissues, we found the variance in genotype explained by LA (the number of African and Asian alleles at the locus) across all 838 genotyped individuals (see Methods). The vast majority of eVariants are not strongly correlated with LA when the entire genotyped population of 838 individuals is considered (Figure 4a).

**Figure 4.**
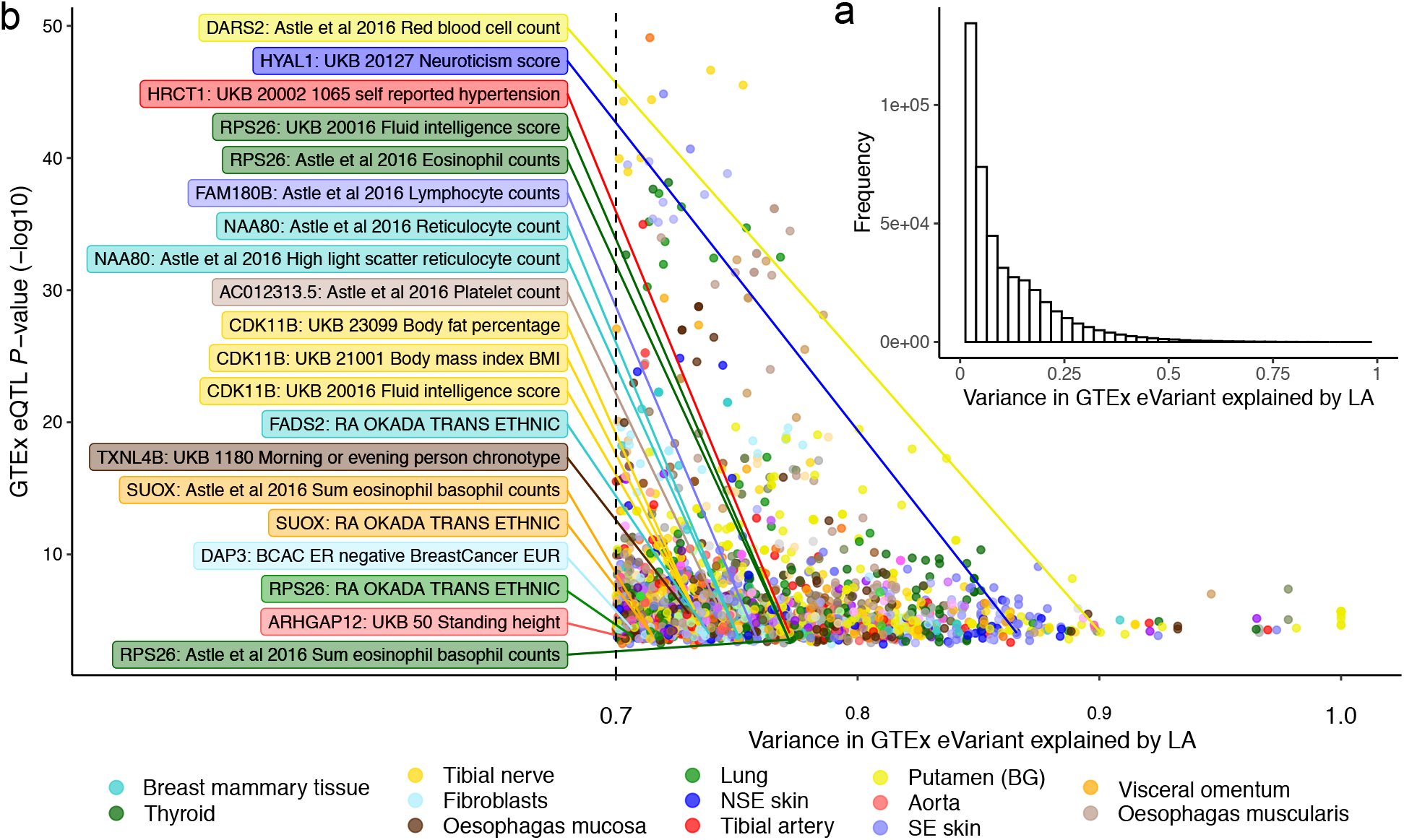
Correlation between genotype and local ancestry in GTEx v8 eVariants. **(a)** The majority of GTEx v8 eVariants are not confounded by local ancestry when all 838 genotyped individuals are considered. **(b)** Local ancestry explains more than 70% of the variance in genotypes for a subset of GTEx v8 eVariants. Unlike **(a)**, **(b)** considers only individuals with matched genotype and gene expression data for each tissue, which reflects the sample used to call these significant associations. eQTLs with posterior probabilities of GWAS colocalization of at least 0.5 are labelled with the eGene and GWAS trait.

However, transcriptome sample sizes within each eQTL tissue are often less than the full sample size (mean 310; standard deviation 171). Therefore, the degree of confounding between a variant’s genotype and LA in the context of eQTL mapping can vary between tissues. To this point, Figure 4b provides the variance in genotype explained by LA for eVariants when only subjects with matched genotype and expression data are included in the regression. Unlike Figure 4a, an eVariant has as many data points as tissues in which it is reported in a significant association. 20 eVariants whose corresponding eGenes have a colocalization probability of greater than 0.5 are also annotated (41). Notably, 19 unique eVariants have proportions of variance explained by LA greater than 0.9 (Table S6, Additional file 2). These variants have large differences in reference allele frequencies between 1000 Genomes populations. For example, one such variant, chr1_1170732_A_G_b38 has reference allele frequencies of 0.993, 0.996, and 0.124 in European, East Asian, and African populations, respectively. A comprehensive list of the 2,556 GTEx v8 significant associations where LA explains more than 70% of the variance in the eVariant genotype are provided in Table S7 (Additional file 2). We expect that functional follow-ups of eQTL/GWAS colocalizations will benefit from crossreferencing with these data.

## Discussion

In this study, we describe population admixture in the GTEx v8 release and assess the impact of ancestry adjustment on eQTLs discovered in an admixed subcohort (117AX). GTEx expands representation from non-European populations, including up to 17% of non-European or admixed individuals. For eQTL mapping, the selection of tissues was limited to those with adequate 117AX samples sizes (>60). We recognize that these relatively small sample sizes will remain an important limitation of multi-population analyses in the GTEx study. Future comparable multi-tissue studies will benefit from increased representation of diverse populations.

The observed trend that GA explains more variance in residual gene expression than LA, on average, agrees with the previous finding that GA explains significantly more heritability of gene expression than LA (42). However, LA can explain a large proportion of variance in GA-corrected gene expression for a subset of genes. Interestingly, a gene whose expression is largely explained by LA, *TBC1D3*, is a highly expanded gene whose copy number is stratified by ancestral population (28,43). Given that copy number expansion is a local phenomenon that has limited effects on global gene expression, population differences in gene copy numbers creates a scenario in which we would expect LA to explain more variance in gene expression than GA. This biological explanation for the differences in *TBC1D3* expression explained by ancestry highlights a specific benefit of considering LA during eQTL mapping.

We decided to include only admixed samples in eQTL mapping on the basis that we would not expect LocalAA to perform any better than GlobalAA in homogeneously European individuals, where the LA covariates are expected to be constant across the majority of the genome. For this same reason, we also excluded homogeneously African (N=14) and Asian (N=9) samples from eQTL calling. However, this does not preclude the use of LocalAA as an ancestry adjustment approach in a cohort with individuals of both homogeneous and heterogeneous ancestry. To this point, Zhong *et al*. reached similar conclusions when comparing LA and GA adjustments in either a strictly African American population or a cohort of mostly European-ancestry individuals with less than 25% African Americans (10).

After performing *cis*-eQTL mapping in six tissues, we observe that LocalAA has a modest improvement in power, consistent with previous observations (10,44). We also observe that most eQTLs agree between LocalAA and GlobalAA; the majority of eGenes that are called uniquely by one ancestry adjustment method are at the threshold of significance. Both of these observations are consistent with previous findings by Zhong *et al*. (10). Further, eGenes called uniquely by GlobalAA are not confounded by LA. Neither do differences in variance in gene expression explained by LA or GA explain these eGenes uniquely called by one method. This, combined with the fact that both methods indicate the same lead eVariant more often than not, even when the association only reaches significance with one method, suggests that eGenes uniquely called by GlobalAA may not in fact driven by confounding with LA. Instead, LocalAA and GlobalAA may have relatively more power for eQTL discovery in different contexts.

To our knowledge, the effects of LA adjustment in eQTL mapping on GWAS colocalization have not previously been explored. We find that neither LocalAA nor GlobalAA in eQTL mapping of six different tissues yields systematically stronger colocalizations across 114 GWAS. In general, stronger colocalization events are captured by both ancestry adjustment methods. Contrary to our expectations, within- and between-continent population differentiation of the LocalAA and GlobalAA eVariants do not differentially impact colocalization probabilities. One colocalization that is significantly stronger with LocalAA than GlobalAA is between the *AP3S2* eQTL in subcutaneous adipose and the DIAGRAM T2D meta-analysis of GWAS in diverse populations, including admixed populations. Possibly LA adjustment in eQTL mapping of an admixed population results in stronger colocalizations with GWAS of admixed individuals, but more work is required to demonstrate this.

Finally, the additional step of LA inference and the incorporation of LA into models for eQTL calling or GWAS makes LocalAA significantly more computationally intensive than GlobalAA. Therefore, a significant improvement of power for discovery or fine-mapping would be required to motivate widespread implementation of LocalAA in large genetic association studies. Several groups recommend that GlobalAA is sufficient to control for type I error during screening for genetic associations, but LocalAA at loci of interest may improve fine-mapping or provide better effect estimates (5,8,9,18). Thus a candidate approach may be taken to adjust for LA only at a subset of loci where LA is expected to improve fine-mapping, which would reduce computational cost and maximize the potential benefit of LA adjustment.

A practical example of this is performing eQTL mapping with GlobalAA and subsequently assessing residual variance explained by LA for discovered eQTLs. To assess this, we post-hoc analyzed GTEx release eVariants to discover 2,556 associations that have large amounts of variance explained by local ancestry (>70%). It remains a challenge to select a threshold for simply excluding QTLs based on the degree of variance explained by local ancestry. We provide this list to enhance future analysis of eQTL/GWAS associations.

## Conclusions

Despite claims of the importance of accounting for LA when performing genetic association studies in admixed populations (15,16), the impact of LocalAA in the context of eQTL mapping has been relatively underexplored. We performed genome-wide LA inference in an admixed subcohort of GTEx v8 and provide these LA calls as a resource to further investigate GTEx data. We then performed *cis*-eQTL mapping in this admixed subcohort to compare GlobalAA and LocalAA ancestry adjustment methods. We observe a modest improvement in power with LocalAA relative to GlobalAA. While both methods yield the same lead eVariant for the majority of eGenes, small subsets of eGenes have different lead eVariants between methods or pass the eQTL significance threshold in only one of the methods. We do not see systematic differences in colocalization probabilities when we perform colocalization between GWAS and eQTLs where the two ancestry adjustments yield different lead eVariants. Interestingly, higher within-continent divergence of lead eVariant allele frequencies is significantly associated with higher GWAS colocalization probabilities for both eQTL ancestry adjustment methods. Finally, we provide a resource of GTEx v8 eVariants that are potentially confounded by LA. Together, these results describe the population structure of admixed individuals in the final GTEx release and demonstrate limited confounding based on local ancestry.

## Methods

### Genotype data

We used GTEx v8 release genotype data (1). Briefly, whole genome sequencing (WGS) was performed for 899 samples from 869 unique GTEx donors, to a median depth of 32x. Alignment to the human reference genome build GRCh38 was performed with BWA-MEM (http://bio-bwa.sourceforge.net). Variants were called with GATK HaplotypeCaller v3.5, and multi-allelic sites were split into biallelic sites using Hail v0.1 (http://hail.is). After performing quality control, the final analysis freeze set contained variant calls from 838 donors. SHAPEIT v2 (45) was used to impute missing calls and phase the sample- and variant-QCed variant call file (VCF).

### Genotype principal component analysis

We used GTEx v8 release genotype principal components (gPCs) (1). gPCs were computed based on the sample- and variant-QCed WGS VCF using EIGENSTRAT (6). PCA was performed on a set of LD-independent variants with a call rate ≥ 99% and MAF ≥ 0.05. LD pruning was performed using PLINK 1.9 (46).

### Gene expression data

We used GTEx v8 release normalized gene expression data; detailed method descriptions can be found in the main GTEx publication (1). RNA sequencing (RNA-seq) was performed at the Broad Institute using the Illumina TruSeq™ RNA sample preparation protocol, which was based on polyA+ selection of mRNA and was not strand-specific. RNA-seq data were aligned to the human reference genome GRCh38/hg38 with STAR v2.5.3a (47). Genelevel expression quantification was performed using RNA-SeQC (48) with a gene annotation available on the GTEx Portal (gencode.v26.GRCh38.genes.gtf). Quantified gene expression (TPM and raw counts) for each tissue was filtered and normalized according to the GTEx eQTL discovery pipeline (https://github.com/broadinstitute/gtex-pipeline/tree/master/qtl). For each of the six tissues in which we chose to perform eQTL mapping, we subsetted normalized gene expression to include only 117AX samples.

### Local ancestry inference

LiftOver (49) was used to convert the phased GTEx v8 whole genome sequencing variant call file (VCF) (dbGaP accession number phs000424.v8) from reference genome Human Build 38 (hg38) to Human Build 37 (hg19) for compatibility with 1000 Genomes and the hg19 HapMap genetic map. The resulting GTEx VCF was filtered to include self-reported African Americans and Asian Americans (103 and 12 individuals, respectively) as well as 25 admixed individuals as identified by the genotype PCA (Figure 1a), resulting in 140 individuals. 1000 Genomes Phase 3 phased VCFs (ftp://ftp-trace.ncbi.nih.gov/1000genomes/ftp/release/20130502) were filtered to include biallelic variants and only individuals in the following populations: Han Chinese in Beijing, China (CHB); Japanese in Tokyo, Japan (JPT); Utah Residents (CEPH) with Northern and Western European Ancestry (CEU); Yoruba in Ibadan, Nigeria (YRI); Gambian in Western Divisions in the Gambia (GWD); Mende in Sierra Leone (MSL); Esan in Nigeria (ESN). The intersection of autosomal variants in the resulting GTEx and 1000 Genomes VCFs (N = ~28M) was identified for LA inference. For compatibility with RFMix v1.5.4, variant positions were converted from base pairs to centimorgans (https://github.com/joepickrell/1000-genomes-genetic-maps) using the HapMap hg19 genetic map (ftp://ftp.ncbi.nlm.nih.gov/hapmap/recombination/2011-01_phaseII_B37).

RFMix v1.5.4 (https://sites.google.com/site/rfmixlocalancestryinference/) was run in PopPhased mode with the additional --forward-backward option. All other parameters were set to the default values. The 1000 Genomes populations were used as reference panels for European (EUR), Asian (ASN), and African (AFR) populations as follows: EUR (CEU, N=99); ASN (CHB, JPT, N=207); AFR (YRI, GWD, MSL, ESN, N=405). This generated posterior probabilities for the assignment of each phased allele to each of the three reference populations (EUR, AFR, ASN). An allele was assigned to a reference population only if the posterior probability was at least 0.9; otherwise, the local ancestry was indicated as “unknown”. For each individual, consecutive phased alleles with the same LA assignment were collapsed into BED files of haplotype blocks with the same LA (Table S2, Additional file 2). These BED files were then used to calculate global ancestry fractions per individual. Scripts used to collapse LA into BED files and calculate global ancestry fractions are available at https://github.com/armartin/ancestry_pipeline.

Of the 140 GTEx v8 individuals whose LA was inferred, 117 individuals with less than 90% global ancestry in a single population (among EUR, AFR, and ASN) were defined as admixed and retained for downstream analyses. This cohort is referred to as 117AX in this paper. VCFtools (50) was used to filter the hg19 GTEx VCF down to variants with a minor allele count (MAC) of at least 10 in 117AX. For the remaining 8,088,666 variants, the LA BED files (Table S2, Additional file 2) were used to count the number of EUR, AFR, ASN, and unknown alleles at each SNP within 117AX. These allele counts were used as LA covariates in eQTL mapping with LocalAA.

### Variance in gene expression explained by ancestry

We adapted an existing approach (26) to quantify variance in gene expression explained independently by LA or GA. For each expressed gene in each tissue, we performed two-step regressions to quantify variance explained by LA (or GA) in gene expression residualized by GA (or LA). First, we regressed out the effects of one type of ancestry on gene expression using the following multiple linear regression, where *γ_i_* is the effect of ancestry covariate *a_i_*, on gene expression *g*, and *e_g_* is the residual:

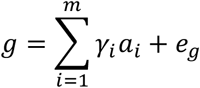

*m*is five for GA (five genotype PCs) and two for LA (numbers of alleles assigned to African or Asian ancestry at the gene’s transcription start site). Then, we quantified variance in *e_g_*explained by the other type of ancestry (*a*\*) by taking the coefficient of determination from the following linear regression:

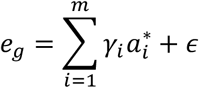

This process was performed for both LA and GA. All regressions were performed with the *lm()* function in R.

### *cis*-eQTL mapping with LocalAA and GlobalAA

Genome-wide *cis*-eQTL mapping in 117AX was performed in six GTEx v8 tissues: Subcutaneous adipose (subc. adipose), tibial artery, lung, skeletal muscle, tibial nerve, and not-sun-exposed suprapubic skin (NSE skin). All methods in this section were performed independently for each tissue. Normalized gene expression files filtered to include only 117AX samples were used to calculate 15 hidden confounders with PEER (51) according to the GTEx eQTL discovery pipeline (https://github.com/broadinstitute/gtex-pipeline/tree/master/qtl). Additional sample-level covariates, including gPCs, WGS sequencing platform (HiSeq 2000 or HiSeq X), WGS library construction protocol (PCR-based or PCR-free), and donor sex were extracted from GTEx v8 release covariate files.

We assumed an additive genetic effect on gene expression and fit the following linear model for each gene-variant pair (gene *g*, variant *v*):

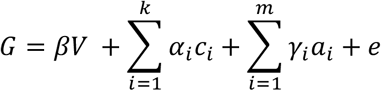

where *G* is expression of gene *g* across 117AX samples in the given tissue, *V* is the number of alternate alleles at variant *v*, coded as 0, 1, or 2; *β* is the effect of the alternate allele of variant *v* on gene *g* expression; *α_i_* is the effect of the technical or biological covariate *c_i_* on gene *g* expression, including donor sex, sequencing platform, library construction protocol, and fifteen hidden confounders; *γ_i_* is the effect of ancestry covariate *a_i_* on gene *g* expression; and *e* is the residual. Any of the 8,088,666 filtered variants within a megabase of the transcription start site of a gene were tested for an association with that gene’s expression. The significance of an association was taken to be the two-sided *P*-value corresponding to the t-statistic of the *β* coefficient estimate. All regressions were performed with the *lm()* function in R.

For each gene-variant pair, two iterations of this regression were performed: one to adjust for global ancestry (GlobalAA), in which case each *a_i_* is one of the first five genotype principal components (gPCs); and one to correct for local ancestry (LocalAA), in which case there are two ancestry covariates, coded as the number of alleles at variant V assigned to African and Asian populations, respectively. gPCs were not included as covariates in the LocalAA model. For LocalAA, samples with any number of alleles with unknown ancestry for the given variant were excluded; the covariates matrix was necessarily reconstructed for each variant tested. This is unlike GlobalAA, where the GA covariates are also sample-level covariates and can be reused for every association test.

After eQTL mapping was completed, the most significant (lead) eVariant (or eVariants, in the case of tied *P*-values) was identified for each gene, independently for the two ancestry adjustment methods. A nominal *P*-value cutoff of 1e-6 was applied to identify significant associations. LD (R^2^) was calculated between single pairs of GlobalAA and LocalAA lead eVariants for each eGene using PLINK (46); an eGene was defined as having different lead eVariants between the two ancestry adjustment methods if 1) there was no intersection between the two sets of lead eVariants and 2) the LD between the tested pair of GlobalAA and LocalAA lead eVariants was less than 1.0.

### Variance in GTEx eVariant genotype explained by local ancestry

In order to identify potential confounding by LA in GTEx v8 eQTLs, we first needed LA calls for all 838 individuals with both WGS and RNA-seq data (1). The remaining 698 individuals for which we did not perform LA inference have self-reported European ancestry and cluster tightly together in gPC space (Figure 1a). Therefore, we approximated LA in these 698 individuals to two European alleles at all tested loci. Then LA covariates for this analysis were the union of computationally inferred LA in 140 admixed or non-European individuals and approximated LA in the remaining 698 homogeneously European individuals.

We calculated the variance explained by LA in the genotype of each eVariant implicated in reported GTEx v8 eQTLs. The following linear model was fit for each eVariant:

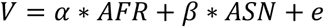

where *V* is the genotype vector (number of minor alleles), and *AFR* and *ASN* are the two LA covariate vectors, representing the number of alleles assigned to African and Asian populations, respectively. The resulting coefficient of determination of each regression was recorded. We did this in two settings: 1) for the set of unique eVariants across all GTEx v8 eQTLs, where genotypes and LA for all 838 individuals were included in the regression (Figure 4a), and 2) for all eVariants within each tissue, with samples subsetted to those with matched gene expression in the given tissue (Figure 4b). 1) provides a global picture of the degree of correlation between eVariant genotypes and LA while 2) reflects the actual samples used to call eQTLs in each tissue. For 2), we also intersected GTEx v8 eQTLs with GTEx v8 GWAS colocalization results (see below) to identify loci with high posterior probabilities of colocalization between eQTLs and GWAS (PP4 > 0.5) associated with eVariants whose genotypes are highly correlated with LA (R^2^ > 0.7).

### Imputation of GWAS summary statistics

Harmonization and imputation of previously published GWAS are described in detail by (41) and (1). Briefly, summary statistics were harmonized and lifted over to hg38; an in-house implementation of BLUP (Best Linear Unbiased Prediction) (52,53) was used to impute z-scores for those variants reported in GTEx without matching data in the GWAS summary statistics.

### Colocalization between eQTL and GWAS signals

Colocalization between 117AX eQTL signals and GWAS signals was performed with COLOC (29). For each of the six tissues in which eQTL mapping was performed, the eQTL summary statistics were subsetted to include only associations with eGenes that had different lead eVariants between the two ancestry adjustment methods (LocalAA and GlobalAA) in any of the six tissues. Colocalization was performed between each subset of eQTL summary statistics (one for each ancestry adjustment method and tissue; twelve in total) and each of the 114 imputed GWAS summary statistics. Nominal *P*-value cutoffs of 1e-5 and 1e-4 were applied to the GWAS and eQTL summary statistics, respectively, in order to select genes that were eligible for colocalization analyses. The reference genome, used to calculate eQTL effect allele frequencies, was the same GTEx VCF used for eQTL mapping in 117AX. eQTL effect sizes and standard errors and GWAS *P*-values were used for colocalization. The seed variant was set to be the GWAS lead variant to have identical seed variants for corresponding colocalizations in the two ancestry adjustment methods; variants within a 500kb window of the seed variant were considered. The COLOC “type” parameter in the *coloc.abf()* function was set to “quant” (rather than case-control “cc”) for all GWAS due to the unavailability of ratios of cases to controls.

Figure 4b references colocalizations identified by an independent analysis of the 114 imputed GWAS and eQTLs reported in the GTEx v8 release (41). Briefly, COLOC was used to perform colocalization with variants in the *cis*-window of each gene with at least one eVariant (*cis*-eQTL per-tissue q-value < 0.05). For binary GWAS traits, case proportion and ‘cc’ trait type parameters were used. For continuous GWAS traits, sample size and ‘quant’ trait type parameters were used. In both cases, imputed or calculated z-scores were used as effect coefficients in Bayes factor calculations. enloc enrichment estimates (54) were used to define data-based priors for COLOC in a consistent manner with other GTEx companion papers (41).

### F_ST_ analyses

All F_ST_ were obtained using the VCFtools calculation of Weir and Cockerham F_ST_ (55). For each variant, 20 within-continent F_ST_ were calculated: 10 pairwise calculations between the five 1000 Genomes European populations (CEU, TSI, FIN, GBR, IBS), and 10 pairwise calculations between the five 1000 Genomes continental African populations (YRI, LWK, GWD, MSL, ESN). The maximum of these 20 within-continent F_ST_ were recorded as F_ST,within_ for each variant. These subpopulations were also combined into European and African populations to calculate F_ST_ between Africa and Europe (F_ST,between_) for each variant.

Linear regression was used to assess the effect of F_ST,between_ and F_ST,within_ of lead eVariants on GWAS colocalization probability. First, we considered results from LocalAA analyses. We identified the highest LocalAA colocalization probability per eGene per tissue (PP4_LocalAA_). Then we matched each eGene with F_ST,between_ and F_ST,within_ for the corresponding LocalAA lead eVariants (F_ST,between,LocalAA_ and F_ST,within,LocalAA_, respectively). If there was more than one LocalAA lead eVariant for an eGene, F_ST,between,LocalAA_ and F_ST,within,LocalAA_ were defined as the maximum of the values within the tied lead eVariants. Then we fit the following model: PP4_LocalAA_ ~ F_ST,between,LocalAA_ + F_ST,within,LocalAA_. We repeated this process for GlobalAA analyses. Coefficient estimates for F_ST,between,LocalAA_, F_ST,within,LocalAA_, F_ST,between,GlobalAA_, and F_ST,within,GlobalAA_ are provided in Figure 3d. Loci tested for colocalization were matched between the two regressions.

## Declarations

### Ethics approval and consent to participate

While deceased individuals do not require consent for research, GTEx consent is described in detail in (56).

### Consent for publication

Not applicable.

### Availability of data and materials

GTEx v8 release gene expression data and *cis*-eQTL call sets are available through the GTEx Portal, https://gtexportal.org. Details of the availability of the 114 GWAS summary statistics used for colocalization are provided in (41). Analysis scripts are available at https://github.com/nicolerg/gtex-admixture-la.

### Authors’ contributions

N.R.G. performed all analyses involving 117AX, local ancestry, or F_ST_; wrote the first draft of the manuscript; and prepared all figures, tables, and supplementary information. M.G. developed the wrapper pipeline that N.R.G. used to perform colocalization. M.G., A.R. provided feedback about colocalization analyses. B.B., S.M. provided advice about statistical analyses. A.R.M., M.L.A., S.M. contributed to discussions about ancestry inference algorithms and software. F.A. generated the genotype principal components used as a proxy for global ancestry. F.A., K.G.A. generated the expression read counts that were then normalized by F.A. and used for *cis*-eQTL mapping. F.A., K.G.A. generated the GTEx v8 release *cis*-eQTL call sets. A.B., R.B., H.K.I. generated the harmonized and imputed summary statistics for 114 GWAS used for colocalization. Y.P., A.B., F.H., C.D.B., X.W., H.K.I. performed colocalization between GTEx v8 eQTLs and the imputed GWAS summary statistics. Y.P., M.L.A, B.B., M.G. helped revise the manuscript. T.L. provided critical feedback. S.B.M. provided guidance for analyses and helped write and edit the manuscript. All authors read and approved the final manuscript.

## Acknowledgements

We thank Jonathan Pritchard for helpful conversations. We thank Joe Pickrell for his publicly available code. We thank Carlos Bustamante, Brian K. Maples, and Christopher DeBoever for the development and maintenance of RFMix.

## Additional information

**Additional file 1 (.pdf):**

Supplementary figures.

**Additional file 2 (.xlsx):**

Supplementary tables.

