## Supplementary figures for "Impact of admixture and ancestry on eQTL analysis and GWAS colocalization in GTEx"

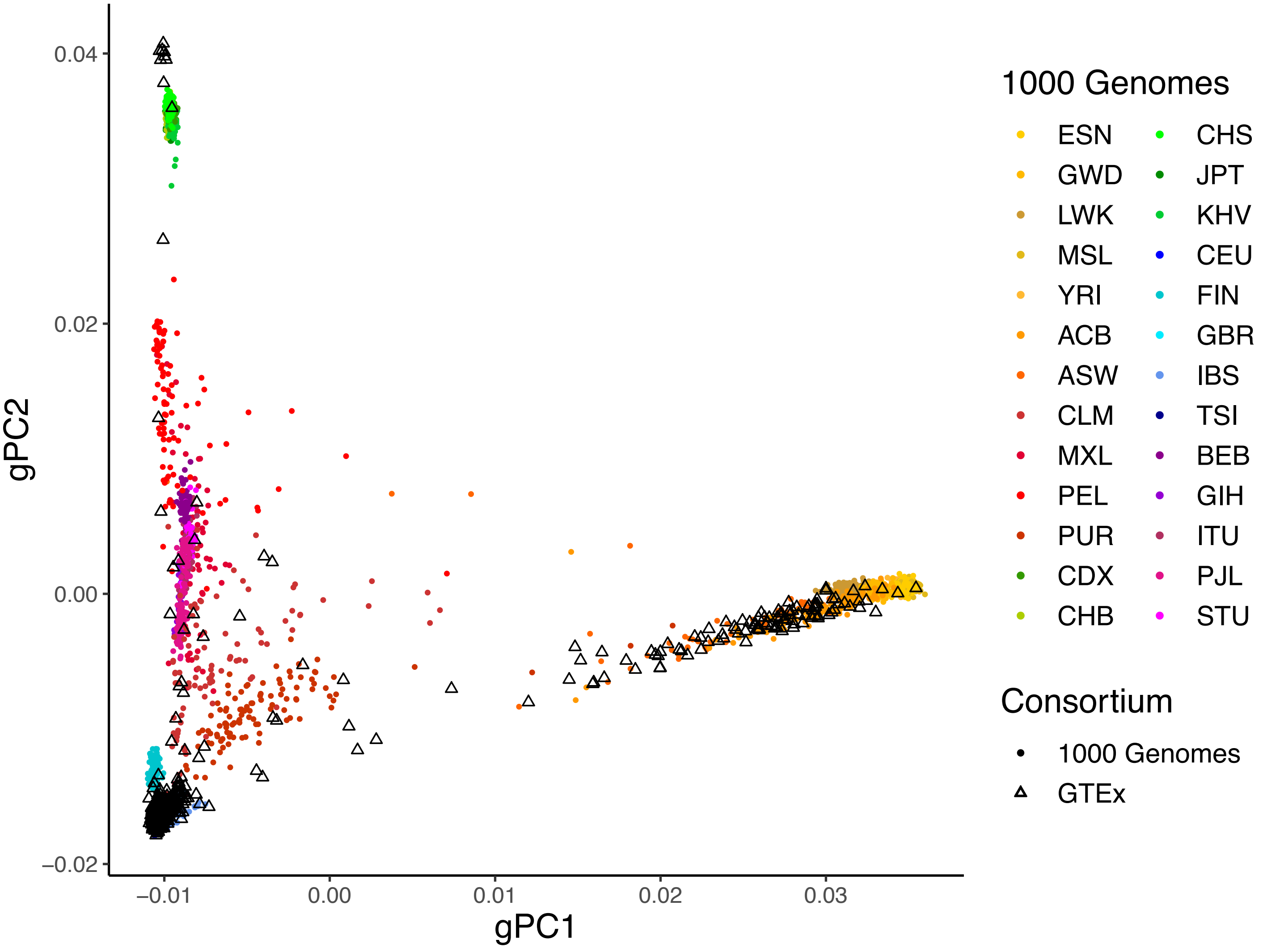


**Figure S1. Spatial distribution of GTEx v8 subjects among 1000 Genomes populations.**
Genotype principal component analysis (gPCA) was performed with combined GTEx v8 and 1000 Genomes genotype data. As in Figure 1a;d, gPC1 is correlated with African ancestry; gPC2 is correlated with Asian ancestry. gPCA was performed using the *snpgdsVCF2GDS()* and *snpgdsPCA()* functions in the SNPRelate R package.

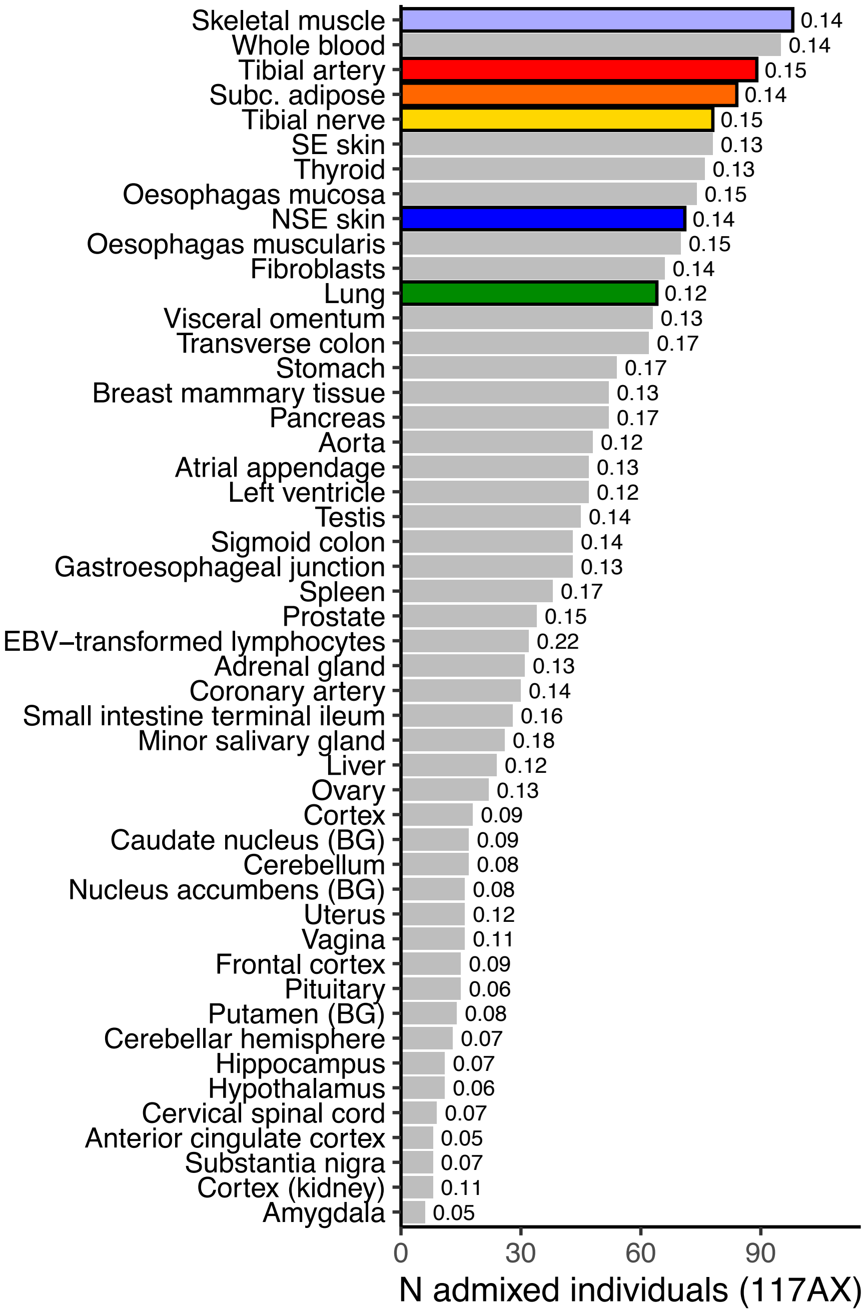


**Figure S2. 117AX sample sizes vary across GTEx v8 tissues.**
The fraction of samples within each tissue that correspond to 117AX samples are indicated to the right of each bar. 117AX represent 0.14 of the total 838 genotyped individuals in GTEx v8. The six tissues selected for eQTL mapping in 117AX are colored.

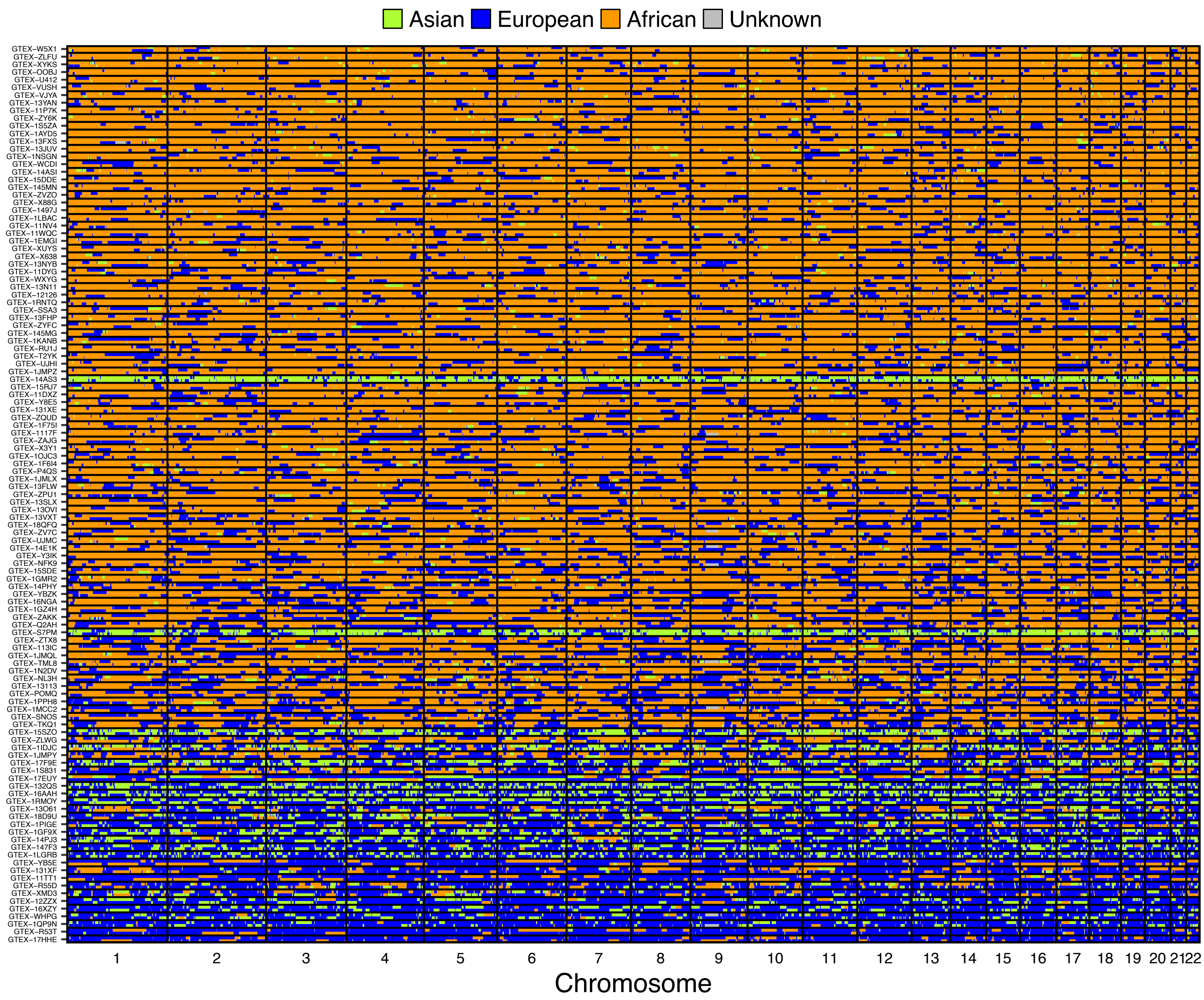


**Figure S3. Genome-wide local ancestry in 117AX.**
Each row provides a visual representation of the local ancestry calls in a haplotype of one individual across autosomal chromosomes (rows). Haplotypes are paired by individual, labelled by GTEx subject ID. Individuals are ordered from top to bottom with increasing amounts of European admixture.

a

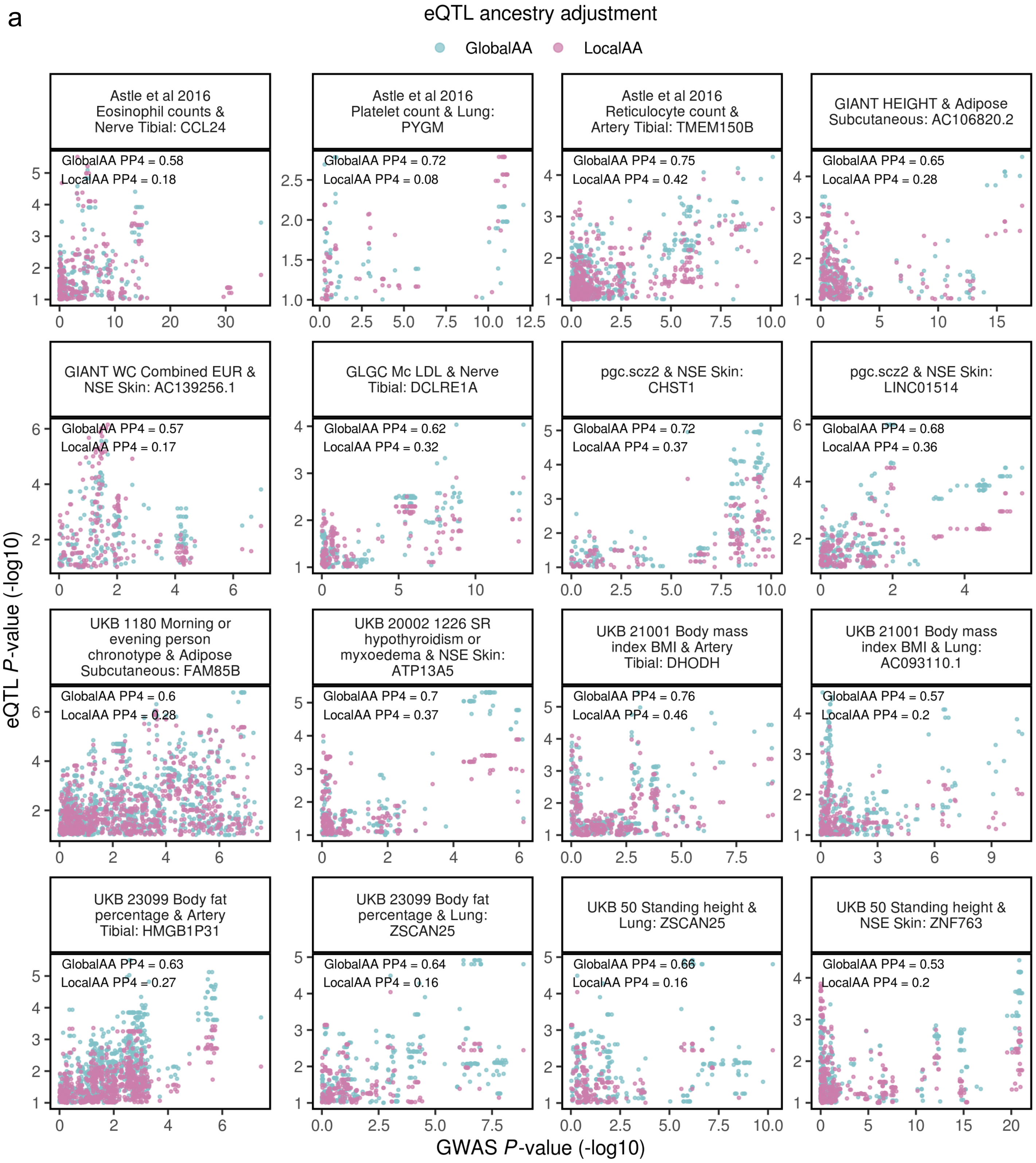
eQTL ancestry adjustment
GlobalAA
LocalAA
Astle et al 2016 Eosinophil counts & Nerve Tibial: CCL24
GlobalAA PP4 = 0.58
LocalAA PP4 = 0.18
Astle et al 2016 Platelet count & Lung: PYGM
GlobalAA PP4 = 0.72
LocalAA PP4 = 0.08
Astle et al 2016 Reticulocyte count & Artery Tibial: TMEM150B
GlobalAA PP4 = 0.75
LocalAA PP4 = 0.42
GIANT HEIGHT & Adipose Subcutaneous: AC106820.2
GlobalAA PP4 = 0.65
LocalAA PP4 = 0.28
GIANT WC Combined EUR & NSE Skin: AC139256.1
GlobalAA PP4 = 0.57
LocalAA PP4 = 0.17
GLGC Mc LDL & Nerve Tibial: DCLRE1A
GlobalAA PP4 = 0.62
LocalAA PP4 = 0.32
pgc.scz2 & NSE Skin: CHST1
GlobalAA PP4 = 0.72
LocalAA PP4 = 0.37
pgc.scz2 & NSE Skin: LINC01514
GlobalAA PP4 = 0.68
LocalAA PP4 = 0.36
UKB 1180 Morning or evening person chronotype & Adipose Subcutaneous: FAM85B
GlobalAA PP4 = 0.6
LocalAA PP4 = 0.28
UKB 20002 1226 SR hypothyroidism or myxoedema & NSE Skin: ATP13A5
GlobalAA PP4 = 0.7
LocalAA PP4 = 0.37
UKB 21001 Body mass index BMI & Artery Tibial: DHODH
GlobalAA PP4 = 0.76
LocalAA PP4 = 0.46
UKB 21001 Body mass index BMI & Lung: AC093110.1
GlobalAA PP4 = 0.57
LocalAA PP4 = 0.2
UKB 23099 Body fat percentage & Artery Tibial: HMGB1P31
GlobalAA PP4 = 0.63
LocalAA PP4 = 0.27
UKB 23099 Body fat percentage & Lung: ZSCAN25
GlobalAA PP4 = 0.64
LocalAA PP4 = 0.16
UKB 50 Standing height & Lung: ZSCAN25
GlobalAA PP4 = 0.66
LocalAA PP4 = 0.16
UKB 50 Standing height & NSE Skin: ZNF763
GlobalAA PP4 = 0.53
LocalAA PP4 = 0.2
eQTL P-value (-log10)
GWAS P-value (-log10)

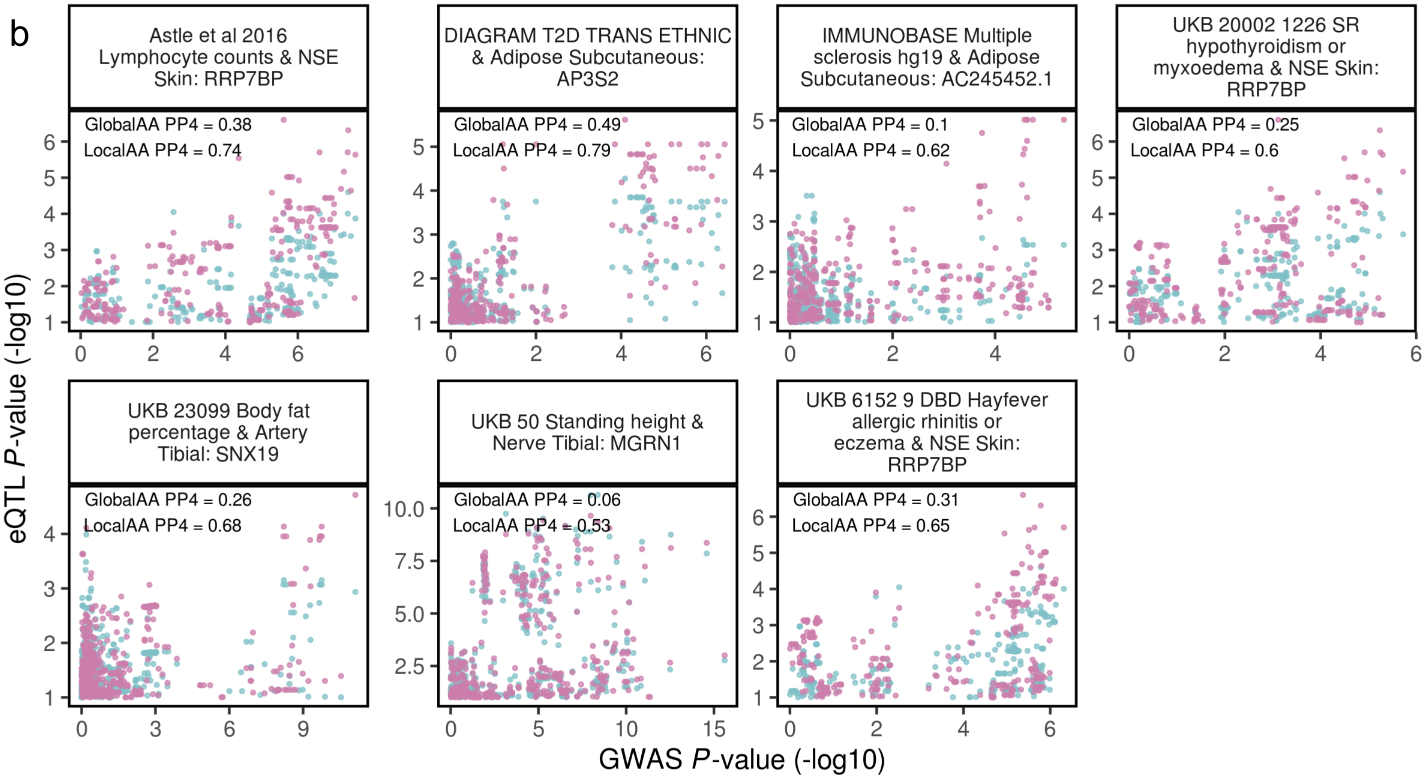


**Figure S4. Loci with stronger GWAS colocalizations using one eQTL ancestry adjustment method.** A colocalization is considered stronger in Method A than Method B if: 1) the colocalization probability (PP4) is greater than 0.5 only in Method A; and 2) Method A PP4 is at least 0.3 greater than Method B PP4. **(a)** 16 colocalizations are stronger with GlobalAA. **(b)** Seven colocalizations are stronger with LocalAA. In general, the shape of the eQTL signals between the two ancestry adjustment methods are similar, but one method has an overall stronger signal. SR = self-reported; DBD = diagnosed by doctor.
